# Hosting sea anemones at the Perhentian reefs of Malaysia: population descriptives and associations with live coral cover

**DOI:** 10.1101/2021.01.22.427756

**Authors:** Melissa Versteeg, Alanah Campbell, Hidayah Halid

## Abstract

Around the Perhentian Islands, coral reefs have been undergoing degradation, as is reported by coral reef monitoring programmes. Current coral reef surveys around the Perhentian Islands do not specifically monitor hosting sea anemone populations, nor do they include investigation of how sea anemone abundance correlates to live coral cover on reef sites. As sea anemones can compete with corals for suitable substrate, nutrients, and light availability, the current study was designed to explore hosting sea anemone abundance and distribution patterns around the Perhentian Islands, as well as assess the presence of significant correlations between sea anemone abundance and live coral cover. Two sites with hosting sea anemone populations were assessed, and data was collected for sea anemone species, formation type, hosting status, and resident Amphiprion species. Additionally, live coral cover estimates were calculated and tested for associations between coral and sea anemone abundance. In total, 403 hosting sea anemone formations were analysed. Statistical analyses revealed that at the research site Village Reef, sea anemones that were actively hosting were larger, and more often encountered in clustered formations. In addition, sea anemone cover was significantly negatively correlated to live coral cover. At research site Teluk Keke, actively hosting sea anemones were also larger, but no other tests revealed significant results at this site. The current study offers a first population analysis of hosting sea anemone assemblages around the Perhentian Islands and provides a preliminary exploration of the associations between hosting sea anemone presence and live coral cover on these reefs.

## INTRODUCTION

On the Perhentian reefs of Malaysia, coral abundance and coral health has been subject to decline, with decreased live coral cover (LCC) reported at longitudinally assessed sites (Reef Check Malaysia, 2007–2019). In 2017, reef assessments conducted by the Perhentian Marine Research Station confirmed this downward trend in coral reef integrity. An analysis of 41 unique sites around the Perhentian Islands indicated a live coral cover average of 27.00 % (SD= 14.000) (Perhentian Marine Research Station, unpublished data). These estimates shift overall reef health at the Perhentian coral reefs from a general classification of ‘fair’ to ‘poor’.

Monitoring of coral reefs around the Perhentian Islands is achieved predominantly through citizen science programmes, which apply simple but effective survey methods using volunteers (; Reef Check Malaysia, 2010; Hunter, Alabri & van Inge, 2013). These approaches collect relevant reef data to calculate reef integrity values, which in turn provide valuable insight for marine park zone designation (Hunter, Alabri & van Inge, 2013; Lau et al., 2019). As determinants of reef integrity, surveys observe various bio-indicators theorised to be related to reef health (Hodgson & Stepath, 1999; Reef Check Malaysia, 2007–2019). Although these methods are valuable and both cost-and time-efficient, they can overlook competition dynamics on the reefs, subsequently introducing risk for misinterpretation, misinformation, and ill-informed management decisions (Wood & Dipper, 2008; Norström et al., 2009; Tun et al., 2013; D’Angelo & Wiedermann, 2014; Tkachenko & Britayev, 2016).

An example of a reef health indicator with more complex mechanisms than are captured in volunteer monitoring programmes, regards nutrient indicator (macro)algae (NIA) as a determinant of coral competition on the reef (Littler & Littler, 2007). In theory, the recording of macroalgae abundance indirectly gauges dissolved nutrient levels, which in turn is negatively associated with coral survival (Littler & Littler, 2013). However, using algae cover to pinpoint eutrophication effects can introduce flaws (Harris, 2015). High macroalgal abundance does not necessarily indicate elevated levels of dissolved nutrients as some macroalgae species can thrive independent of nutrient levels (Harris, 2015). High macroalgal presence may in fact be associated with top-down effects such as overfishing (Norström et al., 2009). More so, not all types of nutrient indicator algae found on the reef detract from coral growth in the same fashion (Littler & Littler, 2007; 2013; Harris, 2015), thus requiring careful interpretation. Another reef health indicator regards sea anemones *(Actiniaria),* which can capitalise off of collapse or imbalance events on coral reefs (Chen & Dai, 2004; Tkachenko et al., 2007; Liu et al., 2009; Tkachenko & Britayev, 2016), but which also associate with live coral in healthy reef settings (Liu et al., 2009). As such, inspecting the abundance of relevant reef species in more detail could overcome inaccuracy pitfalls by presenting a complementary assessment of coral reef competitors and their population dynamics, which is the aim of this research study.

The reef assessments currently used around the Perhentian Islands monitor sea anemone abundance collectively with tunicates, hydroids, and corallimorphs (Reef Check Malaysia, 2019). However, local fishermen have expressed a notable increase in hosting sea anemone abundance, with *Heteractis magnifica* displaying substantial aggregated beds around certain reef regions. Though associated with live coral, increased sea anemone abundance has been found to negatively influence coral planula recruitment and impacts coral recovery rates (Liu et al., 2015; Tkachenko & Britayev, 2016). As such, intensified monitoring of hosting sea anemones is valuable and relevant to better understanding the Perhentian reef dynamics. Furthermore, focussing on hosting sea anemone abundance patterns around these reefs offers exploration of whether these hosting sea anemones are significantly associated with live coral abundance on the coral reefs of the Perhentian Islands. As such, the current study set out to explore relationships between sea anemone presence and live coral cover around the Perhentian Islands, in addition to surveying sea anemone populations to establish a baseline measure for the Perhentian reefs.

Like corals, sea anemones have strict environmental requirements due to their dependency on algal symbionts (Allen, 1975; Fautin & Allen, 1997; Allen et al., 2003), restricting their dominant habitats to the photic zone. Also similar to corals, sea anemones have tentacles with nematocysts for defence, plankton capture, and opportunistic predation (Fautin, 1991). Compared to corals though, sea anemones depend on zooxanthellae to a lesser degree, as they obtain relatively more nutrients though feeding on zooplankton and detritus (Godinot & Chadwick, 2009; Liu et al., 2009). They acquire the bulk of their nutritional needs through zooxanthellic photosynthetic symbionts, in addition to having a capacity for nutrient absorption from the water column through skin tissue (West, de Burgh & Jeal, 1977). Sea anemones also require specific elements for growth including ammonia, phosphate, nitrogen and sulphur (Davies, 1988; Godinot & Chadwick, 2009).

Sea anemones are described as direct coral competitors, and their elevated presence has been reported following outbreaks on newly colonised reefs (Chen & Dai, 2004; Kuguru et al., 2004; Tkachenko & Britayev, 2016). Sea anemone abundance is also reported to be positively influenced by dissolved nutrient levels (Liu et al., 2009; 2015), with suitable environments allowing sea anemone aggregation into extensive beds (Fautin & Allen, 1997; Brolund et al., 2004). Under favourable settings, sea anemones can outcompete stony corals for attachment substrates (Liu et al., 2009). Given sea anemones’ longevity, their potential for year-round asexual reproduction (Fautin & Allen, 1997; Holbrook & Schmitt, 2005), and their fast rate of growth, under positive conditions sea anemones may quickly increase their presence at reef habitats that were previously coral dominated.

Ten sea anemone species have evolved the capacity to host symbiotic anemonefish *(Amphiprion)* (Fautin & Allen 1997). Papua New Guinea is the only current location known to house all species of sea anemones with hosting capacity. For the remainder of the Indo-West Pacific region, prevalence tends to include half of all sea anemone species with hosting capacity (Fautin & Allen, 1997). Around the Perhentian Islands, seven hosting sea anemone species are currently located, including *Heteractis magnifica, Heteractis crispa, Heteractis aurora, Entacmaea quadricolor, Stichodactyla gigantea, Stichodactyla haddoni* and *Stichodactyla mertensii.*

Sea anemones with hosting capacity have the ability to recycle nutrients from waste excreted by symbiotic fish (Holbrook & Schmitt, 2005; Godinot & Chadwick, 2009; Roopin & Chadwick, 2009; Szcezebak, 2013). In fact, sea anemones that successfully host ectosymbionts such as anemonefish have greater concentrations of zooxanthellae, which positively affects growth (Holbrook & Schmitt, 2005). When sea anemones are actively hosting, growth rates have been reported to increase threefold compared to their not actively hosting counterparts. Holbrook & Schmitt’s research also revealed that actively hosting sea anemones have significantly higher asexual reproductive rates than sea anemones without active hosting status, which has been suggested as a driving mechanism for aggregates of identical individuals (Sebens, 1983). The symbiotic relationship between sea anemones and anemonefish also provides benefits at night (Szczebak, et al., 2013). Anemonefish influence oxygen levels of the host sea anemones by altering flow rates around the host tissue during night time. Thus, the symbiotic relationship that hosting sea anemones can maintain with resident anemonefish offers benefits that can facilitate growth, formations, and abundance on coral reefs.

The current study investigated hosting sea anemone distributions at two Perhentian reef sites. Furthermore, this study sought to conduct a preliminary exploration of the associations between sea anemone abundance and live coral presence. The following research questions were formulated: (1) What are the hosting sea anemone species distributions, size estimates, active hosting indicators, formation types, and distribution patterns at Village Reef and Teluk Keke? (2) Are there significant differences in sea anemone size between actively hosting sea anemones versus not actively hosting sea anemones? (3) Are there differences in formation types based on the hosting status of sea anemones? And, (4) are there significant associations between hosting sea anemone presence and live coral cover at the Perhentian reef sites?

## MATERIALS & METHODS

Data was collected at Village Reef (central coordinates: 5°53’39.05” N, 102°43’37.61” E) and Teluk Keke (central coordinates:5°53’14.0316’’N, 102°44’20.9004’’E). Village Reef is also locally referred to as ‘Nemo’ in acknowledgement of its high abundance of hosting sea anemones and anemonefish. It lies on the intertidal zone off the southeast of Perhentian Kecil (**Fig. 1a**). Teluk Keke (**Fig.1b**) is located to the West of Perhentian Besar, and its reef contains rocky areas in combination with sheltered regions of shallow reef.

**Fig. 1.**
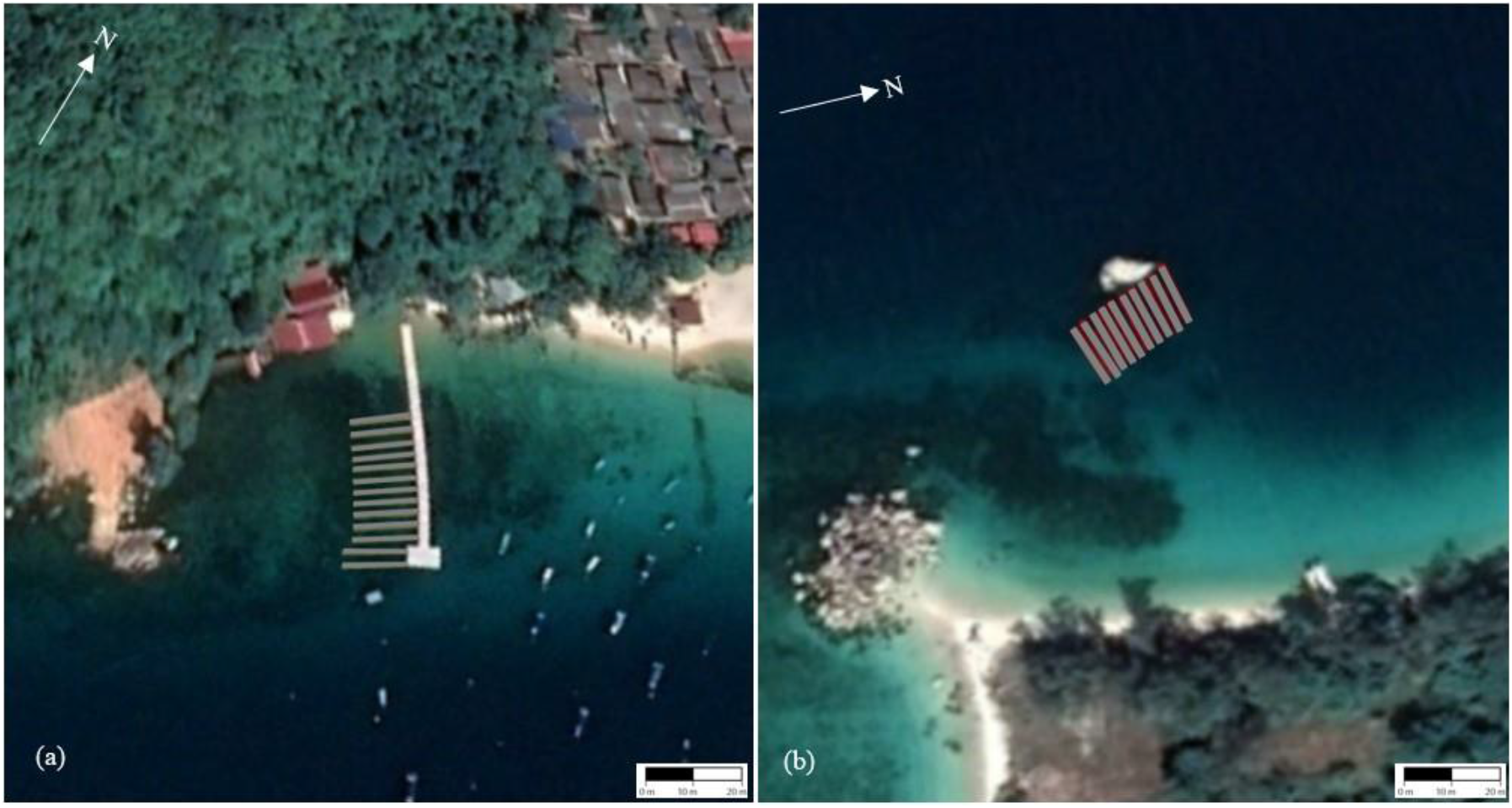
Survey sites Village Reef, at Pulau Perhentian Kecil (a) and Teluk Keke on Pulau Perhentian Besar (b), with depiction of transects within the research sites. *Note.* Image source: Google 2020, CNES/Airbus, 1 cm: 20 m. At Village Reef, the three shallowest transects initially included in the survey site where discarded due to a lack of hosting sea anemone presence.

Between August 5^th^ 2020 and August 20^th^, 2020, SCUBA was used to study hosting sea anemone populations as well as measure coral cover at the two research sites. At site regions too shallow for SCUBA (depth <2.0m), data was collected using freediving techniques. Within the boundaries of survey area Village Reef, a total of ten 20 meter transects were laid out in parallel using a 225° southwest bearing, as well as spatial referencing from a stable landmark. At Teluk Keke, ten 20 meter transects were also laid out in parallel, using a 270° northwest bearing. At Teluk Keke, a partially exposed rock made for a natural landmark for additional spatial referencing. The distance between parallel transects was set at 4 meters to allow optimal observation whilst mitigating inflated counts caused by overlap. Upon laying of the transect, two trained research divers regressed along the line, taking a two-meter perpendicular width and they recorded all relevant study information.

When encountering a sea anemone, a long and short axis measurement of the oral disc was taken using a tailor’s tape. In addition, relevant spatial mapping measures were taken including transect identifiers and transect distance readings. Sea anemone species were visually identified, the formation type was recorded (Allen, 1975; Fautin & Allen, 1997; Allen et al., 2003), hosting status was determined (Fautin & Allen, 1997; Holbrook & Schmitt, 2005), and any resident anemonefish were visually identified for species identification (Fautin & Allen, 1997; Allen et al., 2003; Wood & Aw, 2017). When experiencing ambiguity, video footage was collected to allow cross referencing ex situ. In classifying formations of clusters of sea anemones, individuals were assessed as forming a cluster if a fully expanded individual’s tentacles could touch a neighbouring sea anemone (Sebens, 1983; Brolund et al., 2004).

As formulas to calculate area coverage assume full coverage between the elliptical long and short axis (Hirose, 1985), clusters which did not fully cover the substrate, or clusters which did not assume an elliptical shape, were adjusted for by recording area cover estimates. Furthermore, site rugosity measurements were taken to account for site complexity in subsequent sea anemone cover estimates (Knudby & LeDrew, 2007). To calculate area cover percentages of the sampled sea anemones, cover estimates were divided by transect segment area, which was calculated at 20 m^2^ excluding site complexity adjustments (4-meter width x 5-meter length intervals along the 20-meter transect line, for a total area per transect of 80 m^2^). To calculate live coral cover, the substrate directly underneath the same transect line was visually identified at 50 cm intervals, at a total of 40 points per transect line (Manuputty Djuwariah, 2009). Hard and soft coral data points were subsequently extracted to inform LCC percentage estimates.

All data collection sessions took place between 0830 hours and 1159 hours, and visibility during data collection had to be over five meters as a prerequisite to diving. An interobserver analysis (Hartmann, 1977) revealed an overall recording and identification accuracy of 96,70 %. All statistical analyses were run using IBM SPSS (Statistical Package for the Social Sciences) for Windows, version 27.0.

## RESULTS

At Village Reef, several hosting sea anemone species could be identified, including *Stichodactyla gigantea, Stichodactyla mertensii,* and most notably *Heteractis magnifica* (**Table 1**). As for Teluk Keke, three species of hosting sea anemone were recorded: *Heteractis magnifica*, *Entacmaea quadricolor,* and *Stichodactyla mertensii*. At Teluk Keke *Heteractis magnifica* also demonstrated higher abundance compared to other species (**Table 1**).

**Table 1.**
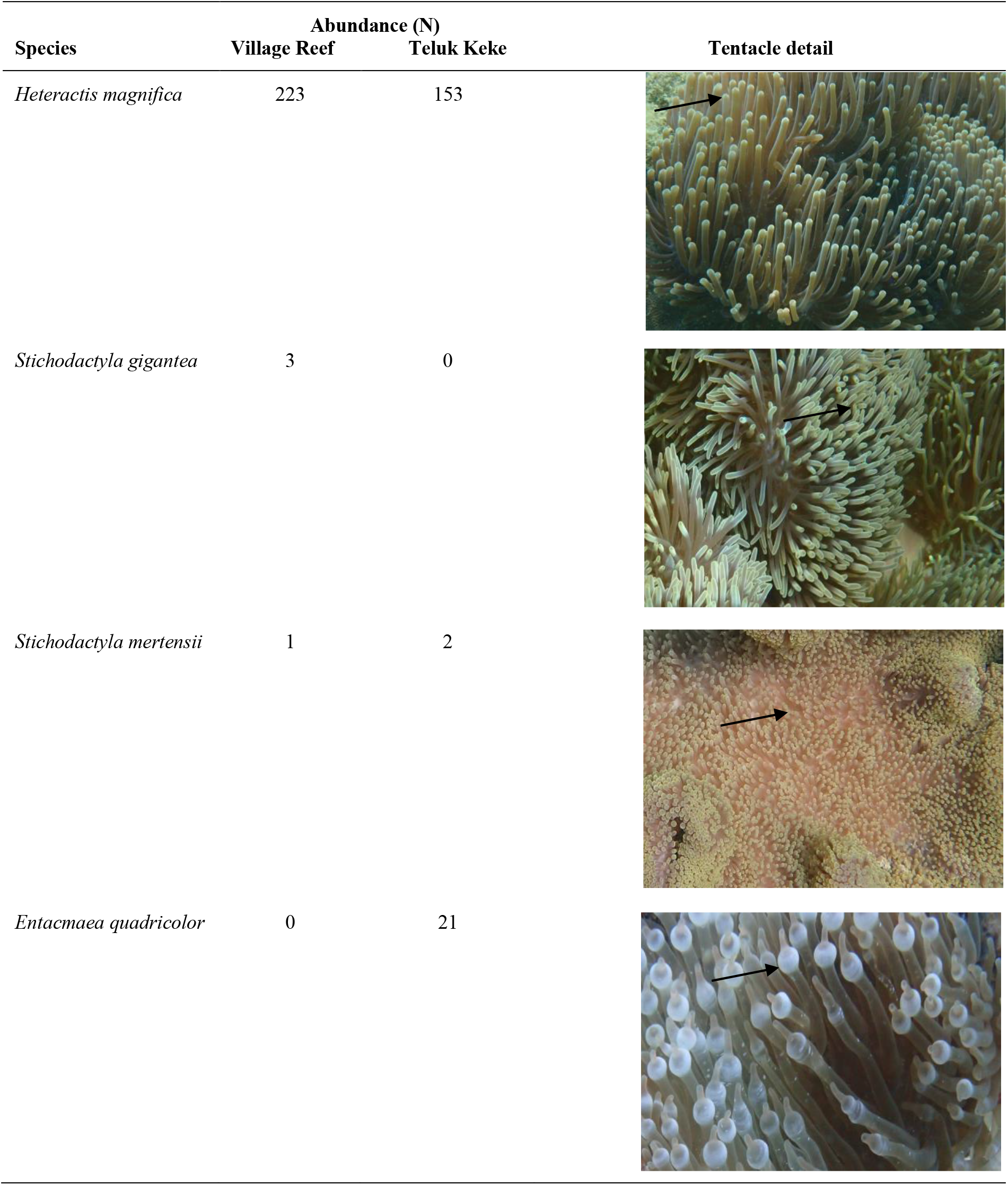
Hosting sea anemone species located at Village Reef and Teluk Keke, including abundance and tentacle detail.

| Species | Abundance (N) |  | Tentacle detail |
| --- | --- | --- | --- |
|  | Village Reef | Teluk Keke |  |
| *Heteractis magnifica* | 223 | 153 |  |
| *Stichodactyla gigantea* | 3 | 0 |  |
| *Stichodactyla mertensii* | 1 | 2 |  |
| *Entacmaea quadricolor* | 0 | 21 |  |

At Village Reef, a total of 227 sea anemone formations were identified and analysed. *Heteractis magnifica* was the most dominant sea anemone species, with 98.24 % presence (N= 223). Furthermore, three specimens of *Stichodactyla gigantea* were identified, and one *Stichodactyla mertensii* specimen was recorded (**Table 2**). The average size of all studied sea anemones was 0.129 m^2^ (SD= 0.195 m^2^, MIN= 0.002 m^2^, MAX= 1.891 m^2^). The average size of just *Heteractis magnifica* sea anemones was 0.130 m^2^ (SD= 0.197 m^2^, MIN= 0.002 m^2^, MAX= 1.891 m^2^). The total cover of hosting sea anemones at Village Reef was 29.32 m^2^ and the total calculated cover pertaining solely to *Heteractis magnifica* was 29.05 m^2^.

**Table 2.** Population descriptors for the hosting sea anemones at Village Reef, including size and cover estimates for hosting status, formation types, and sea anemone species.

| **Village Reef** |  |  |  |  |
| --- | --- | --- | --- | --- |
|  | **N** | **Mean size (m<sup>2</sup>)** | **SD (m<sup>2</sup>)** | **Area Cover (m<sup>2</sup>)** |
| *Heteractis magnifica* | 223 | 0.130 | 0.197 | 29.050 |
| *Stichodactyla gigantea* | 3 | 0.063 | 0.017 | 0.189 |
| *Stichodactyla mertensii* | 1 | 0.083 | n.a. | 0.083 |
| Total | 227 | 0.129 | 0.195 | 29.322 |
| Hosting Status |  |  |  |  |
| Active Hosting | 176 | 0.157 | 0.209 | 27.369 |
| Not active hosting | 51 | 0.038 | 0.096 | 1.953 |
| Formation Type |  |  |  |  |
| Solitary | 117 | 0.038 | 0.031 | 4.471 |
| Cluster <5 | 62 | 0.116 | 0.099 | 7.177 |
| Cluster 6–10 | 26 | 0.240 | 0.098 | 6.252 |
| Cluster 11–15 | 13 | 0.340 | 0.127 | 4.420 |
| Cluster 16+ | 9 | 0.778 | 0.478 | 7.002 |
| *Heteractis magnifica*<br>(N= 223) |  |  |  |  |
| Hosting Status |  |  |  |  |
| Active Hosting | 173 | 0.157 | 0.210 | 27.179 |
| Not active hosting | 50 | 0.037 | 0.096 | 1.871 |
| Formation Type |  |  |  |  |
| Solitary | 113 | 0.037 | 0.030 | 4.199 |
| Cluster <5 | 62 | 0.116 | 0.099 | 7.177 |
| Cluster 6–10 | 26 | 0.240 | 0.098 | 6.252 |
| Cluster 11–15 | 13 | 0.340 | 0.127 | 4.420 |
| Cluster 16+ | 9 | 0.778 | 0.478 | 7.002 |

For all hosting sea anemones at Village reef, 77.53 % were actively hosting *Amphiprion spp.* at the time of analysis (N= 176). Of *Heteractis magnifica,* 77.58 % were actively hosting (N= 173). Of the actively hosting sea anemones surveyed at Village Reef, 84.09 % hosted *Amphiprion ocellaris* symbionts (N= 148), 15.34 % were found to host *Amphiprion perideraion* (N= 16), and one formation hosted both *Amphiprion ocellaris* and *Amphiprion perideraion* at 0.57 %. As for formations (**Table 2**), for all sea anemone species, 51.41 % were solitary formations, with the remainder clustered in formation (**Table 2**). Regarding *Heteractis magnifica,* 50.67 % of the sample contained solitary formations, with the remainder present in clustered formation (**Table 2**).

The hosting sea anemone cover estimates were also calculated per transect (**Table 3**) to allow analysis of the relationship between live coral cover and sea anemone abundance. The average percentage cover of all sea anemones for the ten transects was 3.67 % per transect (MIN= 0.56 %, MAX= 11.96 %), with an average live coral cover of 39.00 % (MIN= 17.50 %, MAX= 60.0 %). Other descriptives are further presented in **Table 3**.

**Table 3.**
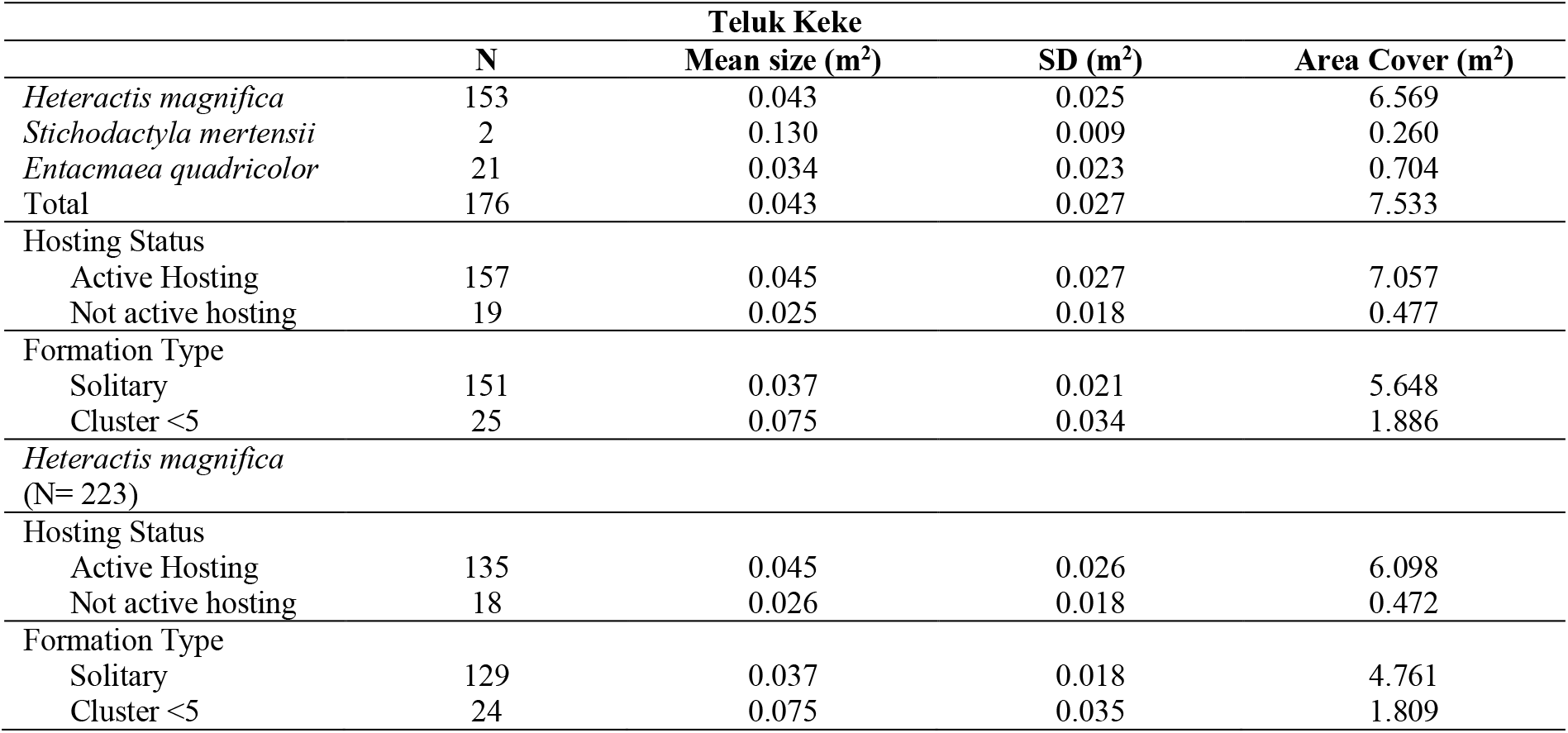
Population descriptors for the hosting sea anemones at Teluk Keke, including size and cover estimates for hosting status, formation types, and sea anemone species.

At Teluk Keke, a total of 176 sea anemones formations were identified and analysed. Here, *Heteractis magnifica* was also the most dominant species, with 86.93 % presence (N= 153). *Entacmaea quadricolor* species had the second highest abundance levels, at 11.93 % (N= 21). Just two specimens of *Stichodactyla mertensii* were recorded (**Table 1**), representing 1.14 % of the total sample. The average size of all studied sea anemones was 0.043 m^2^ (SD= 0.027 m^2^, MIN= 0.005 m^2^, MAX= 0.161 m^2^). The average size of *Heteractis magnifica* was also 0.043 m^2^ (SD= 0.025 m^2^, MIN= 0.008 m^2^, MAX= 0.161 m^2^). The total cover of hosting sea anemones at Teluk Keke was 7.533 m^2^ and the total calculated cover pertaining solely to *Heteractis magnifica* was 6.569 m^2^.

For all hosting sea anemones at Teluk Keke, 89.20 % were actively hosting *Amphiprion spp.* (N= 157) (**Table 4**). For *Heteractis magnifica* only, 88.24 % were actively hosting (N= 135). Of these actively hosting sea anemones at Teluk Keke, 76,43 % hosted *Amphiprion ocellaris* symbionts (N= 120), and 10.19 % were found to host *Amphiprion perideraion* (N= 16). Furthermore, *Amphiprion frenatus* was found to reside on 12.74 % of actively hosting sea anemones (N= 20) and one sea anemone was actively hosting *Amphiprion clarkii,* at a percentage of 0.64 %.

**Table 4.** Area and percentage cover for hosting sea anemones and LCC percentages at Village Reef and Teluk Keke.

| Hosting Sea Anemone Descriptives |  |  |  |  |  |  | LCC |  |
| --- | --- | --- | --- | --- | --- | --- | --- | --- |
| Village Reef (VR) |  |  |  | Teluk Keke (TK) |  |  | VR | TK |
| Transect | Formations (N) | Cover (m<sup>2</sup>) | Cover (%) | Formations (N) | Cover (m<sup>2</sup>) | Cover (%) | Cover (%) | Cover (%) |
| 1 | 52 | 7.504 | 9.38 | 1 | 0.036 | 0.05 | 20.00 | 7.50 |
| 2 | 75 | 7.226 | 9.03 | 3 | 0.192 | 0.24 | 35.00 | 25.00 |
| 3 | 47 | 9.568 | 11.96 | 13 | 0.477 | 0.60 | 17.50 | 27.50 |
| 4 | 29 | 3.025 | 3.78 | 19 | 0.653 | 0.82 | 22.50 | 15.00 |
| 5 | 10 | 1.203 | 1.50 | 39 | 1.717 | 2.15 | 45.00 | 17.50 |
| 6 | 6 | 0.353 | 0.44 | 35 | 1.470 | 1.84 | 45.00 | 15.00 |
| 7 | 3 | 0.167 | 0.21 | 37 | 1.734 | 2.17 | 60.00 | 20.00 |
| 8 | 1 | 0.045 | 0.06 | 17 | 0.755 | 0.94 | 50.00 | 32.50 |
| 9 | 2 | 0.144 | 0.18 | 11 | 0.470 | 0.59 | 45.00 | 27.50 |
| 10 | 2 | 0.086 | 0.11 | 1 | 0.030 | 0.04 | 50.00 | 12.50 |

As for formations (**Table 4**), sea anemones at Teluk Keke were found only in solitary formations or in clusters including less than five individuals. For all sea anemone species, 85.80 % were solitary formations, with the remainder clustered in formations of less than five individuals (**Table 4**). Regarding *Heteractis magnifica,* 84.31 % regarded solitary individuals. Relevant descriptives are further presented in **Table 4**.

The highest sea anemone coverage was localised around the centre of survey site Teluk Keke (**Table 3**). The average LCC at Teluk Keke was 20.00 % (MIN= 7.50 %, MAX= 32.50 %), with an average hosting sea anemone cover of 0.94 % (MIN= 0.04 %, MAX= 2.17 %) per transect.

To control for interspecies differences in size, abundance patterns, and formations (Allen, 1975; Fautin & Allen, 1997; Allen et al., 2003), data pertaining only to *Heteractis magnifica* was extracted to answer the second and third research question. To test for differences in size between the actively hosting sea anemones versus not active hosting sea anemones, a Mann-Whitney U test was used as data was non-normal (Village Reef: Kolmogorov-Smirnov=.527, p<.001; Teluk Keke: Kolmogorov-Smirnov=.127, p < .001). Results revealed a significant difference in sea anemone size at Village Reef (U= 7625.000, p < .001,SE= 401.826, N= 223) for actively versus not actively hosting *Heteractis magnifica*, where the actively hosting sea anemones are significantly larger (**Table 3**). The Mann Whitney U test for Teluk Keke also revealed a significant difference in size between active and not active hosting status of *Heteractis magnifica* (U= 1849.000, p < .001,SE= 176.589, N= 153), with larger sizes recorded for the actively hosting sea anemones (**Table 3**).

Further statistical testing was conducted to assess whether the hosting status of sea anemones, active versus not active, at Village Reef and Teluk Keke is significantly related to their formation, based on previous research indicating that actively hosting sea anemones engage in higher rates of asexual reproduction (Fautin & Allen, 1997; Holbrook & Schmitt, 2005) which is argued to underlie clustered formations of individuals (Sebens, 1983; Fautin & Allen, 1997; Brolund et al., 2004). As such, Chi-Square tests were conducted to test the relationship between hosting status and cluster formations for *Heteractis magnifica* at both survey sites. Results demonstrate that actively hosting *Heteractis magnifica* were more often encountered in clusters at Village Reef (X^2^(6) = 40.892, p <.001), though results for Teluk Keke were marginally nonsignificant (X^2^(1) = 3.795, p=.051).

Finally, to test whether hosting sea anemone presence significantly correlates with live coral cover, as has been argued in previous research (Liu et al., 2009; Tkachenko & Britayev, 2016), sea anemone cover and live coral cover at both survey sites were analysed using a Spearman’s correlation test (**Table 4** and **Figure 2**). Results of the correlation analysis indicate that at Village Reef, hosting sea anemone cover significantly negatively correlates with live coral cover (Spearman’s rho= -.886, p=.001, N= 10). Higher levels of sea anemone cover at Village Reef are associated with lower levels of live coral cover. As for Teluk Keke, no significant associations were found between sea anemone presence and live coral cover.

**Fig. 2.**
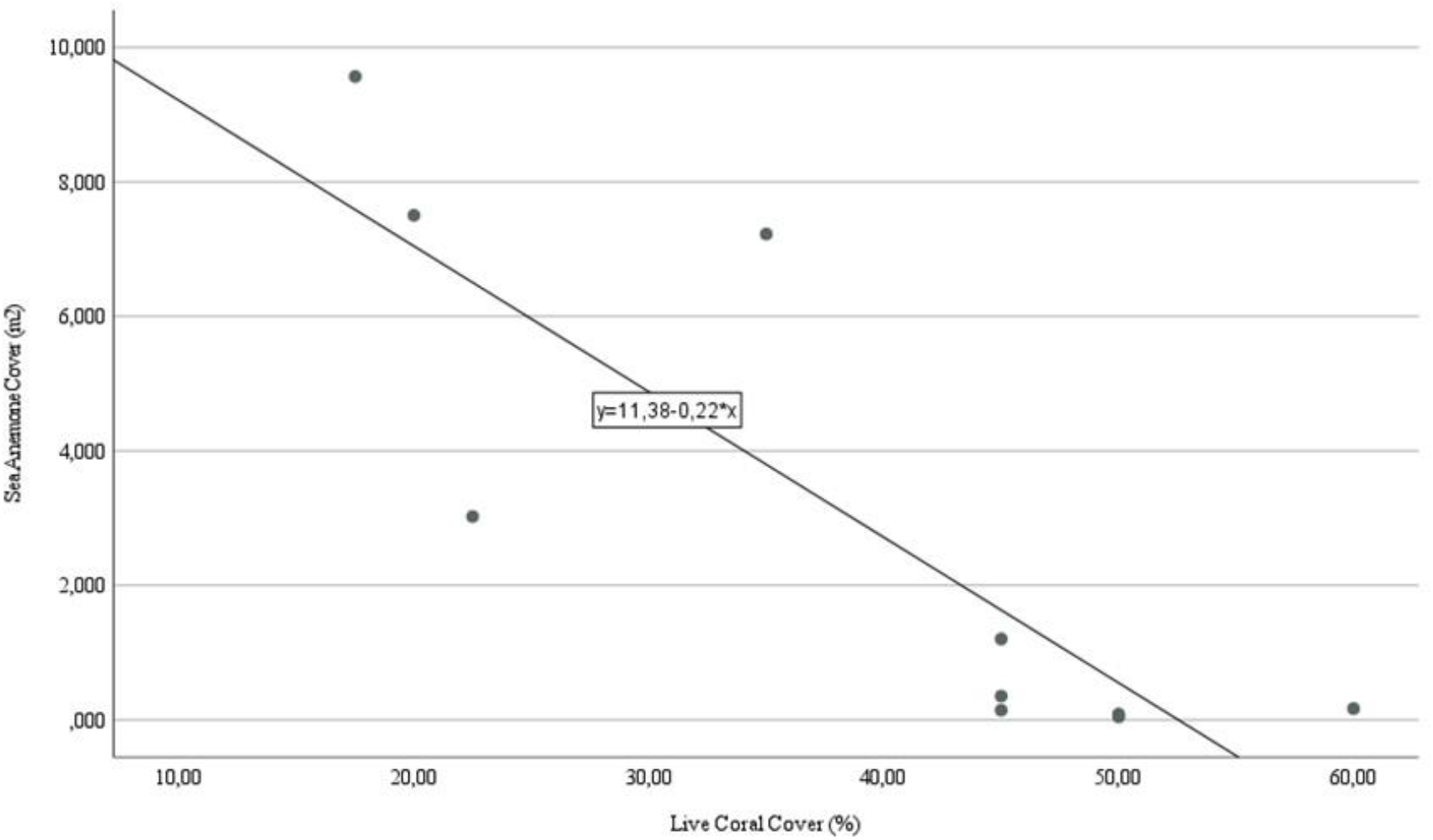
Scatter plot including line of best fit displaying the association between hosting sea anemone cover and live coral cover at Village Reef. *Note.* R^2^= 0.714.

## DISCUSSION

The current study sought to provide preliminary insight on hosting sea anemone assemblages found around the Perhentian islands, including an investigation into their population descriptives and associations with live coral presence on the reefs. Two survey sites were assessed, and data was collected on species distributions, size estimates, hosting status, and formation types. More so, the study wanted to assess whether there were size differences in sea anemones based on hosting status, whether actively hosting sea anemones were more often encountered in clusters, as well as exploring associations between hosting sea anemones and live coral cover.

At Village Reef, hosting sea anemone distributions were more concentrated at the deeper transects of the site. Similar to findings at Teluk Keke, the dominant species regarded *Heteractis magnifica,* although the presence of *Stichodactyla gigantea* was unique to Village Reef. Also unique to Village Reef was the larger presence of clustering formations of hosting sea anemones. At Village Reef hosting sea anemones were larger and more often found in clustered formation. More so, a negative correlation was seen when comparing hosting sea anemone cover to live coral cover. On transects with higher levels of sea anemone cover, live coral cover estimates were generally lower.

The findings related to Village Reef support previous research on the ability of hosting sea anemones to outcompete corals (Liu et al., 2009; Tkachenko & Britayev, 2016). More so, almost half of all surveyed sea anemones at this site were clustered in formation, which has been proposed to indicate increased asexual reproductive success, and is thought to underlie higher levels of sea anemone aggression (Turner et al., 2003; Holbrook & Schmitt, 2005). The sea anemones that were actively hosting *Amphiprion* at Village Reef were also more often encountered in clusters. The sea anemones’ higher ability to absorb waste excreted by resident fish, which in turn stimulates growth and asexual reproductive rates (Holbrook & Schmitt, 2005; Liu et al., 2009; Roopin & Chadwick, 2009; Cleveland, Verde & Lee, 2011) likely drives this finding at Village Reef.

The study outcomes related to Teluk Keke demonstrated both similarities and differences compared to Village Reef. At Teluk Keke, actively hosting sea anemones were also significantly larger, a finding that is in line with previous research (Holbrook & Schmitt, 2005; Liu et al., 2009; Roopin & Chadwick, 2009; Cleveland, Verde & Lee, 2011). In contrast to results from Village Reef, sea anemones at Teluk Keke were not encountered in clustered formation more often. It might be that the specimens at Teluk Keke were still in juvenile stages, as the average size of specimens located at Teluk Keke was smaller, and as juvenile sea anemones are believed not to cluster with the same frequency as adults (Turner et al., 2003).

At Teluk Keke, *Entacmaea quadricolor* specimens were recorded, a species which was not located at Village Reef. More so, the analysis revealed no significant associations with live coral cover at Teluk Keke. It could well be that environmental factors present at Teluk Keke are substantially different from Village Reef, which in turn influences the local population dynamics and microhabitat use on the reef (Chomsky et al., 2004; Dixon et al., 2014). The lack of findings regarding hosting status and formation types could also be the result of reduced statistical power, as only a small number of hosting sea anemones at this site were clustered in formation. With continued monitoring of this site, the new questions that have arisen can be investigated.

### Limitations

Although we aimed to maintain the best standards for scientific rigour, the current study has several limitations. First of all, assessment of hosting status was conducted using in-water direct observation by trained researchers. Though the inter-observer accuracy was high, data collection methods using in-water observations to assess fish behaviours can introduce some disadvantages compared to the use of video recording techniques (Branconi, Wong & Buston, 2019), which may have influenced the accuracy of the hosting status observations as presence of the diver may have impacted resident fish behaviours and visibility.

Second, size estimates for the hosting sea anemones were collected using the oral disc diameter as opposed to the pedal disc diameter. Scientific consensus posits that the pedal disc diameter is preferable, as oral disc measurements are subject to diurnal expansion rates (Allen, 1975). However, the presence of large clustered formations at Village Reef in addition to the high structural complexity found at Teluk Keke drove the decision to measure oral disc diameters, and to estimate cluster sizes using the short and long axis across the aggregated clustered formation. As such, inaccuracies due to expansion or contraction behaviours could have been introduced into the data, although all data collection dives were set to occur in the mornings to control for such effects.

Third, the current study only assessed two Perhentian reefs as a consequence of the novel coronavirus pandemic during the timing of the study. As a result, findings related to Village Reef and Teluk Keke have yet to be compared to other sites around the Perhentian islands, which means that caution should be taken when extrapolating the current findings to other sea anemone populations around the Perhentian Reefs. Fourth and finally, the live coral cover estimates were calculated using a simplified strategy compared to the methods used to estimate hosting sea anemone cover. As such, fewer data points were available for live coral cover estimates, which, should erroneous readings have been present, could have a disproportionate effect on coral estimates. Replication studies should be done to ensure accuracy of the current findings when comparing coral and sea anemone cover.

### Practical implications and future directions

This study provided a first investigation into hosting sea anemone populations around the Perhentian Islands of Malaysia. In line with previous research, the sea anemones that were actively hosting were significantly larger than not actively hosting sea anemones, which provides evidence that these populations are benefiting from the presence of symbiotic anemonefish (Hollbrook & Schmitt, 2005; Godinot & Chadwick, 2009; Liu et al., 2009; Roopin & Chadwick, 2009; Cleveland, Verde & Lee, 2011). Additionally, evidence was presented to indicate that actively hosting sea anemones were also more often found in clustered formation, and associations were found to suggest that, in areas with higher abundance of hosting sea anemones live coral levels were lower, which is in line with prior research in Southeast Asia (Tkachenko & Britayev, 2016).

The findings of the study imply that, at Village Reef, the sea anemone population display growth and reproduction behaviours that are similar to other geographical regions and laboratory settings (Holbrook & Schmitt, 2005; Liu et al., 2009). More so, with the identification of clustered sea anemones within extensive aggregates, the current study supports previous reports on the ability of sea anemones to aggregate in waters around Malaysia (Fautin & Allen, 1997; Allen et al., 2003; Brolund et al., 2004; Wood & Aw, 2017), and extends these findings to include the Perhentian reefs as a location where such aggregates can be found. The current findings are highly relevant as previous studies mention a lack of available data on sea anemone abundance on coral reefs (Norström et al., 2009). By providing a first assessment of hosting sea anemones on the Perhentian reefs, the current study offers baseline population descriptions that can inform population trends in upcoming research.

The study provides several important directions for future research. Regarding the two sites that were included, future research should continue to focus their research efforts on these sites, as longitudinal trends can be studied using the current results as a baseline (e.g. Versteeg, Campbell & Halid, preprint). Furthermore, to allow general population estimates for the Perhentian Islands the amount of research sites should be expanded. Sites without marked hosting sea anemone presence may also be included in future research so that the association between live coral cover and sea anemone abundance can be further explored, in addition to allowing deeper exploration of impacted corals at the genus level (Tkachenko & Britayev, 2016).

Finally, the current research set-up could yield more widespread implications by striving to include abiotic measures in upcoming surveys. Measuring influential factors such as nutrient levels, water temperature, sedimentation, soft coral presence, bleaching events, and algal abundance (Nugues & Roberts, 2003; Chomsky et al., 2004; Wood & Dipper, 2008; Tun et al., 2013) will enhance the potential to provide instrumental insights. By including such factors, results may tap into localised expansive behaviours of sea anemones, algal dynamics can be inspected to asses coral and sea anemone competition dynamics, the sensitivity of corals and sea anemones to bleaching can be examined, and valuable information on the abundance of other implicated benthic invertebrates can be obtained. Collectively, such continued research effort into sea anemone abundance at the Perhentian reefs will help to improve the accuracy of coral reef integrity measures, and it will contribute pertinent information in support of reef management and conservation efforts.

## AKNOWLEDGEMENT

This research and its resultant report were produced with the support of Perhentian Marine Research Station (https://www.marineresearchstation.org/). Many thanks to the team, and specifically to Fuze Ecoteer Cofounder Daniel Quilter. The data collection was conducted at reef sites Village Reef and Teluk Keke (Pulau Perhentian Kecil and Besar, respectively) with the Coral Rehabilitation Project in Perhentian Island permit JTLM 610-4/1/1 Jld 3 (28) and the Strategic Partnership: Reef Care Program permit Prk. ML. 610-3/1 (9), both issued by Malaysian Marine Parks Department Taman Laut Malaysia (http://marinepark.dof.gov.my). The current research was carried out with the aim of informing rehabilitation prospects through habitat composition surveys, in addition to establishing baseline monitoring data on site dynamics. A big thank you is due to everyone who supported the research dives by donating to the fundraiser, by helping with dive logistics, or simply by supporting the work of the Perhentian Marine Research Station. Terima kasih banyak!

## DATA AVAILABILITY STATEMENT

The data presented in this study are available upon request from the corresponding author*. The data are not publicly available due to legal publishing constraints as defined in the regulations inherent to the permits issued by Taman Laut Malaysia.

